# Oncogenic mutant RAS signaling activity is rescaled by the ERK/MAPK pathway

**DOI:** 10.1101/2020.02.17.952093

**Authors:** Taryn E. Gillies, Michael Pargett, Jillian M. Silva, Carolyn Teragawa, Frank McCormick, John G. Albeck

**Affiliations:** Department of Molecular and Cellular Biology, University of California, Davis, CA; UCSF Helen Diller Family Comprehensive Cancer Center, San Francisco, CA; Frederick National Laboratory for Cancer Research, Frederick, MD, USA

## Abstract

Activating mutations in RAS are present in ∼30% of human tumors, and the resulting aberrations in ERK/MAPK signaling play a central role in oncogenesis. However, the form of these signaling changes is uncertain, with activating RAS mutants linked to both increased and decreased ERK activation *in vivo*. Rationally targeting the kinase activity of this pathway requires clarification of the quantitative effects of RAS mutations. Here, we use live-cell imaging in cell lines expressing only one RAS isoform to quantify ERK activity with a new level of accuracy. We find that despite large differences in their biochemical activity, mutant KRAS isoforms within cells have similar ranges of ERK output. We identify roles for pathway-level effects, including variation in feedback strength and feedforward modulation of phosphatase activity, that act to rescale pathway sensitivity independent of expression level, ultimately resisting changes in the dynamic range of ERK activity while preserving responsiveness to growth factor stimuli. Our results reconcile seemingly inconsistent reports within the literature and imply that the initial signaling changes induced by RAS mutations in oncogenesis are subtle.

## Introduction

The RAS GTPases act as molecular switches, alternating between an inactive GDP-bound state and an active GTP-bound state. In the active state, RAS proteins have a greatly increased binding affinity for their effectors (Gremer et al., 2011), which in mammalian cells drive multiple cell growth signaling pathways. The net signaling activity of RAS in the cell represents a balance between two classes of proteins: GTPase-Activating Proteins (GAPs), which inactivate RAS by increasing its GTPase activity, and Guanine nucleotide Exchange Factors (GEFs), which catalyze the dissociation of GDP and return RAS to the active GTP-bound state. Though RAS proteins are considered binary switches on the molecular level, the collective behavior of the thousands of RAS proteins present inside each cell is analog in nature. The relative activity of GAPs vs. GEFs in the cell determines the fraction of RAS molecules in the active state, which in turn regulates the activity of downstream processes.

RAS mutations occur frequently in cancers, especially those of the pancreas, lung, or colon (Fernandez-Medarde and Santos, 2011), and typically have the effect of increasing the signaling output of one of the RAS isoforms. Most oncogenic RAS mutations (85%) occur in the KRAS isoform, with 11% in NRAS and 4% in HRAS (An and Harper, 2018). Across all isoforms, 98% of oncogenic mutations are located at G12, G13 and Q61 (Prior et al., 2012) and render the RAS proteins GAP-insensitive to varying degrees. The net effect of these mutations is to increase the fraction of RAS proteins in the GTP-bound active state, which enhances their binding affinity to effectors, including the RAF kinases (Gremer et al., 2011; Hunter et al., 2015; Smith et al., 2013). RAF initiates a kinase cascade involving MEK and ERK (Fig. 1A), which plays a primary role in tumor development and is a pharmacological target for cancer therapy. ERK phosphorylates hundreds of downstream targets (Yoon and Seger, 2006), many of which are transcription factors controlling cell cycle progression and cell migration.

**Figure 1.**
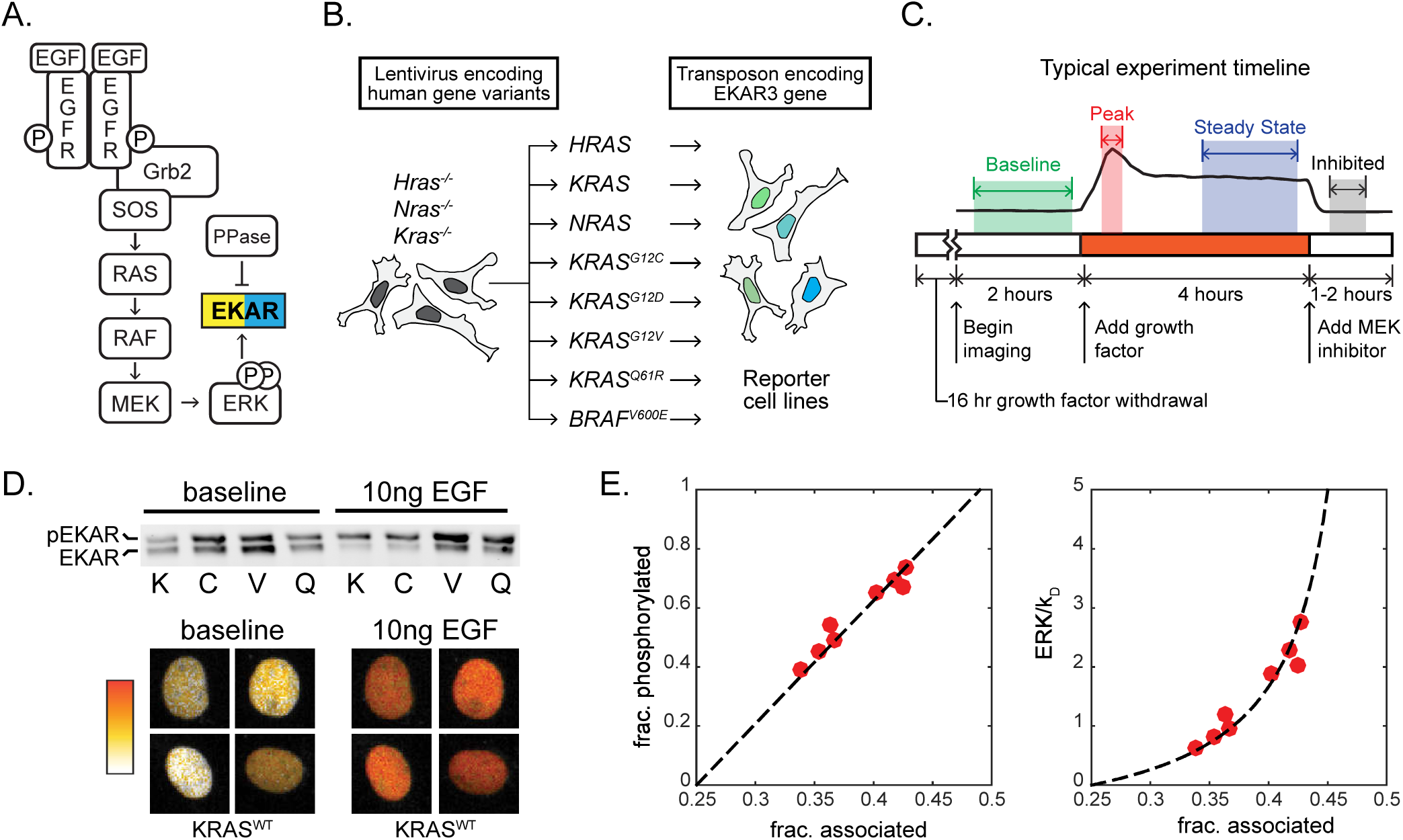
Platform for ERK activity measurement in MEF cell lines expressing a single RAS isoform. A) Schematic of EGF signaling through RAS to ERK, including the EKAR3 sensor. B) Construction scheme for cell lines bearing a single RAS isoform, using H/K/N-RAS knockouts. C) Diagram of the typical experiment timeline. Shaded regions indicate time windows that are averaged for each measurement. D) Sample calibration data for the EKAR3 reporter, consisting of Phos-Tag immunoblot for phospho-EKAR (upper) and live-cell imaging of reporter FRET activity (lower) under matched conditions for 4 cell lines. Four single nuclei from the KRAS^WT^ line are shown before and after stimulus. E) Calibration curves for ERK activity. Fraction of EKAR3 phosphorylated is shown vs. the fraction in the associated conformation by FRET (left). The ERK to phosphatase activity ratio (right) is derived from a model of EKAR3 (see Supplemental Methods). Each red dot represents the mean value from one cell line with or without EGF treatment, from 3 replicate live-cell cultures and 4 replicate western blot cultures.

While RAS mutations are widely thought to initiate tumors by enhancing the activity of RAF/MEK/ERK signaling to drive tumorigenic cellular behaviors, this model is not consistent with all of the data available. A number of observations deviate from this simple linear view of RAS signaling. First, the observed frequency of RAS mutations in cancer does not correlate with the strength of their effect on RAS GTPase activity. Mutations of intermediate strength are most prevalent, and the strongest mutations are found infrequently (Li et al., 2018). Second, the mutational status of RAS is poorly correlated with average levels of active dually phosphorylated ERK (ppERK) both in tumor cell lines (Omerovic et al., 2008; Yeh et al., 2009) and in genetically engineered mouse models. In fact, converting a wild type *Kras* gene to an activating mutant can actually *reduce* average ppERK levels, despite inducing tumor formation (Tuveson et al., 2004). These data contrast with observations that ectopic expression of RAS mutants does drive strong over-activation of ERK, as would be expected from simple amplification by RAF/MEK/ERK (Konishi et al., 2007; Park et al., 2006).

These contradictory observations could arise for various reasons, including feedback in the RAS/ERK pathway (Courtois-Cox et al., 2006), oncogene-induced senescence (Sarkisian et al., 2007), additional mutations, or tissue- or cell type-specific effects (Brandt et al., 2019). However, these possibilities cannot be disentangled without first addressing a major technical limitation inherent in the immunoblots and kinase assays that have been used almost exclusively to date. Because immunoblot measurements are relative, not absolute, it is not typically possible to compare the magnitude of ERK activation across datasets or studies. Furthermore, ERK activation has been shown to be pulsatile and heterogeneous (Albeck et al., 2013; Regot et al., 2014), so the relative differences observed when blotting for active ERK could have multiple biochemical interpretations: more ERK proteins may be active per cell, or cells with active ERK may be more frequent in the population (Birtwistle et al., 2012; Purvis and Lahav, 2013). Increases in the magnitude, frequency, or duration of ERK activation pulses would all yield the same result via immunoblot, though each of these signaling changes would imply different effects on gene expression and warrant different approaches for pathway directed therapy.

To clearly distinguish the forms of ERK activity that result from RAS mutations, we combined live-cell and immunoblot techniques to study a panel of cell lines each expressing only one wild type or mutant isoform of human RAS in an isogenic background. To unequivocally measure ERK activity, we employed a genetically encoded Förster Resonance Energy Transfer (FRET)-based sensor (EKAR3), and calibrated it to deliver a quantitative linear readout of ERK substrate phosphorylation. The live-cell sensor allows measurement of cell-to-cell heterogeneity and signaling dynamics for a complete view of ERK activity in each cell. Complementing live-cell data with immunoblot measurement of RAS/ERK pathway components and computational modeling, we found that ERK activity is strikingly constrained in cells expressing mutant KRAS. When unstimulated, KRAS mutant cells exhibit only moderately elevated ERK activity compared to the wild type, and when stimulated reach peak activity no greater than the wild type. These findings outline a new unified model for how elevated RAS activity is modulated by downstream effectors and for which signaling characteristics may be relevant in cancer.

## Results

### A platform to quantify ERK activity downstream of individual RAS isoforms

To evaluate the cellular signaling capacity of each RAS isoform individually, we utilized a panel of genetically engineered mouse embryonic fibroblasts (MEFs) in which the genes for the three major RAS isoforms (*Hras*, *Kras* and *Nras*) have been functionally deleted and complemented with a single constitutively expressed human cDNA (Drosten et al., 2010) (Fig. 1B). Human proteins expressed are: HRAS, KRAS, NRAS, KRAS^G12C^, KRAS^G12D^, KRAS^G12V^, KRAS^Q61R^, or the oncogenic RAF gene BRAF^V600E^, in which case no RAS isoform is expressed. In these cell lines the signaling behavior of each RAS protein isoform can be characterized in isolation both from other isoforms and from locus-specific variations in transcriptional regulation. To track the resulting signaling activity with high temporal resolution, we transfected each MEF cell line with EKAR3, a live-cell FRET-based ERK activity reporter (Harvey et al., 2008; Sparta et al., 2015) (Fig. 1A). These cell lines were imaged by time-lapse fluorescence microscopy and analyzed with a custom image analysis pipeline (see Methods), typically yielding 100-300 single-cell time series measurements of ERK activity from each replicate of an experimental condition.

To enable accurate comparisons between the single-RAS cell lines, we developed a workflow to make quantitative live-cell measurements of ERK activity (Fig. 1C). Signaling activity in the absence of external stimulation, a condition we term “baseline”, was quantified in cells cultured with neither serum nor growth factors for at least 16 hours prior to imaging. Responses to receptor stimulation were quantified by introducing growth factor after several hours of baseline imaging. As a negative control for ERK reporter measurements, we treated cells with the highly specific MEK inhibitor PD0325901 (MEKi), which rapidly inhibited the EKAR3 signal in all cell lines. In all live-cell experiments, the 100nM MEKi treatment was applied just prior to ending the experiment; this measures the cell-specific residual EKAR3 signal, accounting for non-specific fluorescence. The signal from the EKAR3 reporter was derived from the intensity ratio of the cyan and yellow fluorescent channels (CFP/YFP) and corrected for background as well as excitation and filter spectra. The corrected EKAR3 signal linearly reflects the fraction of reporter molecules in a FRET conformation, which is in turn linearly related to the fraction of molecules phosphorylated by ERK (Birtwistle et al., 2011). To calibrate, we used Phos-Tag immunoblotting to quantify the fraction of the EKAR3 reporter that is phosphorylated in various samples and conditions (Fig. 1D) and fit these values against the average corrected EKAR3 signal for the same cell lines and conditions. In concert with a mass action model of substrate phosphorylation, this calibration yields a linear measure of the ERK:phosphatase ratio (Fig. 1E), i.e. the concentration of active ERK divided by the concentration of any active phosphatases that dephosphorylate the reporter (see Supplemental Methods for details). As these competing phosphatases also have affinity to endogenous ERK targets, the ERK activity measurement reflects not just the levels of active ERK, but the *net* effect of ERK on its substrates.

To demonstrate the utility of this platform for assessing inhibitor activity, we treated the panel of reporter cells with ARS-853, an inhibitor specific to KRAS^G12C^. Following treatment with ARS-853, ERK activity decreased over the course of 60 minutes in KRAS^G12C^ MEFs, but not in any of the other KRAS cell lines (Fig. 2A). Thus, allele-specific drug responses can be quickly identified and quantified using the reporter cell panel.

**Figure 2.**
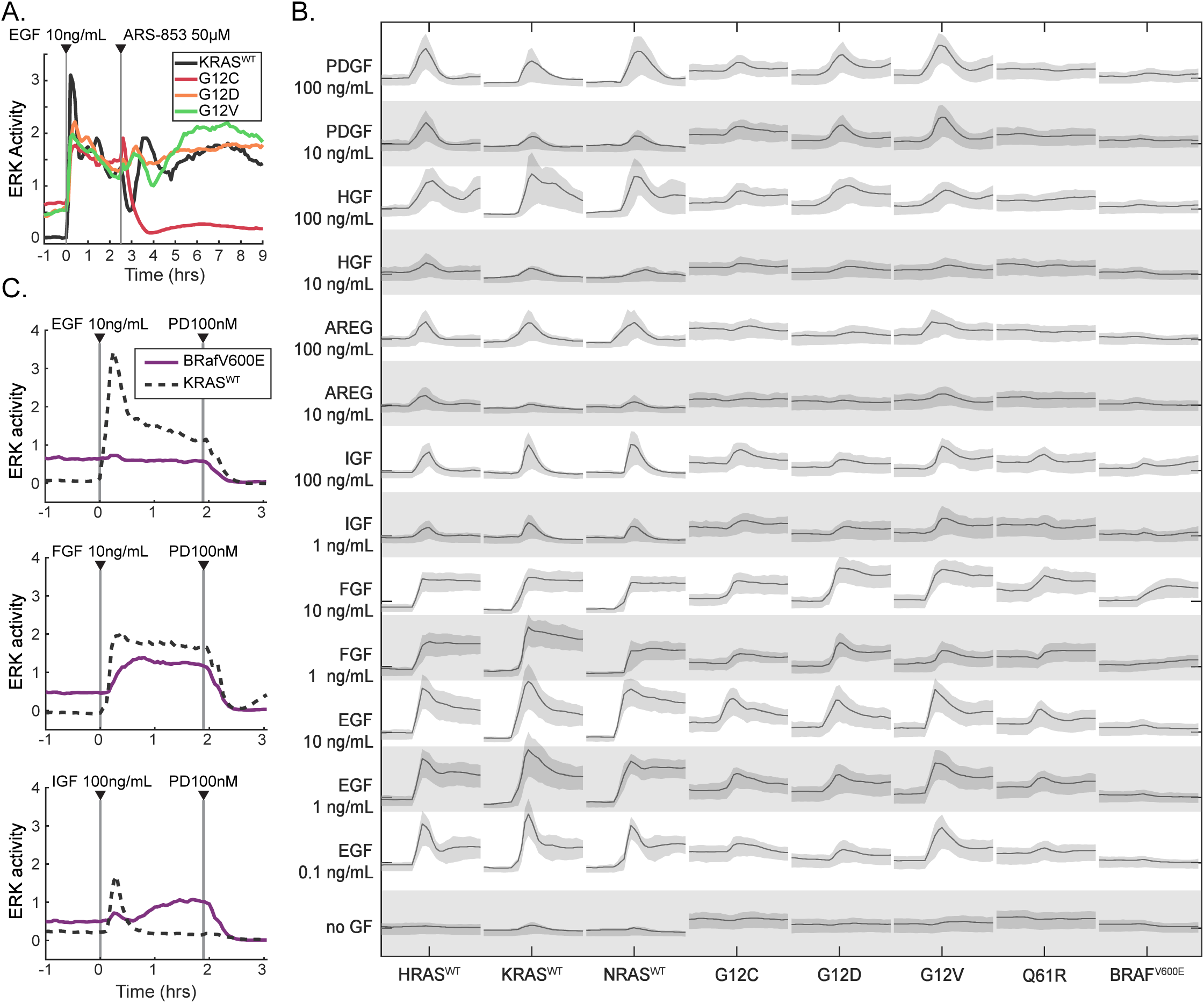
Activity profiles of MEF cell lines expressing a single RAS isoform. A) Demonstration of the system measuring a cell line specific response via ARS-853, a RAS activity inhibitor specific to the KRAS^G12C^ mutant. Traces are median values from a representative experiment. B) Graphical summary of single RAS cell lines (labelled along bottom) stimulated by a panel of growth factors (labelled along left). Each panel of the matrix shows the time series of ERK activity with the indicated growth factor spiked in after beginning imaging. All scales are equal; x-axis: time; y-axis: ERK activity. Lines indicate median of single cell measurements over time, and shaded regions denote the 25^th^ – 75^th^ percentile region, across 3 replicate cultures (6 for no GF). C) Demonstration of RAS-independent activity from ligands other than EGF, evidenced by response in the BRAF^V600E^ cell line lacking H/K/N RAS. Traces are median values from a representative experiment.

We next used our platform to perform a comprehensive survey of the effects of various growth factors on each cell line. We stimulated the MEF cell line panel with 6 growth factors known to activate RAS/ERK signaling: EGF, IGF, FGF, HGF, PDGF, and Amphiregulin. Two to three concentrations of each growth factor were tested across three biological replicates, yielding activity “traces” from approximately 400 cells per condition (Fig. 2B). ERK kinetics differed depending on the growth factor. For example, FGF induced sustained ERK activity without pulsatile behavior, while IGF induced a single ERK activity pulse, approximately 30-40 minutes in duration, immediately following stimulation. By virtue of having no reintroduced RAS genes, the BRAF^V600E^ cell line was not expected to respond to any growth factor stimulus. However, both FGF and IGF induced elevated ERK activity in BRAF^V600E^ cells (Fig. 2C), indicating an ERK response that is not mediated via RAS, which would confound our comparisons of the RAS isoforms. By contrast, EGF induced high amplitude ERK activity in both mutant and wild type RAS cells, without evidence of RAS-independent activity in the BRAF^V600E^ cell line. The remaining growth factors, PDGF, HGF, and Amphiregulin, did not induce activity in BRAF^V600E^ cells, but induced weaker or more transient ERK responses than did EGF in RAS-expressing cells. We therefore focus on EGF for the bulk of our subsequent analysis, because it induces a strong ERK response without RAS-independent effects.

### RAS mutants only moderately elevate ERK activity, and only without stimulation

With quantitative single-cell resolution available, we addressed the question of how the RAS protein isoforms differ in their ERK activity patterns. After growth factor withdrawal, cells were stimulated with either media alone, or media with EGF to a final concentration of 10 ng/mL (Fig. 3A-C). Across all cell lines, EGF stimulation initiated a rapid ERK activity peak ∼15 minutes after stimulation, followed by attenuation over 1.5 – 2 hours to reach a steady state level (Fig. 3B,C), with HRAS^WT^ and NRAS^WT^ cells exhibiting slower attenuation than any of the KRAS isoforms. Responses in single cells were qualitatively similar to the average, though each cell showed variation over time (Fig. 3C). To statistically compare responses across single cells, we decomposed each single-cell ERK trace into parameters: average baseline activity, peak stimulated activity, stimulated amplitude, and average steady state activity 2 hours after stimulation. To compare the tendency for sporadic and time-varying activity, we computed a “volatility” metric as the mean absolute derivative of ERK activity in a single cell, scaled by the mean for that cell. This metric is the time-dependent equivalent of the coefficient of variation; low volatility indicates flatter more consistent activity, while higher volatility indicates more pulses, or changes in activity level over time.

**Figure 3.**
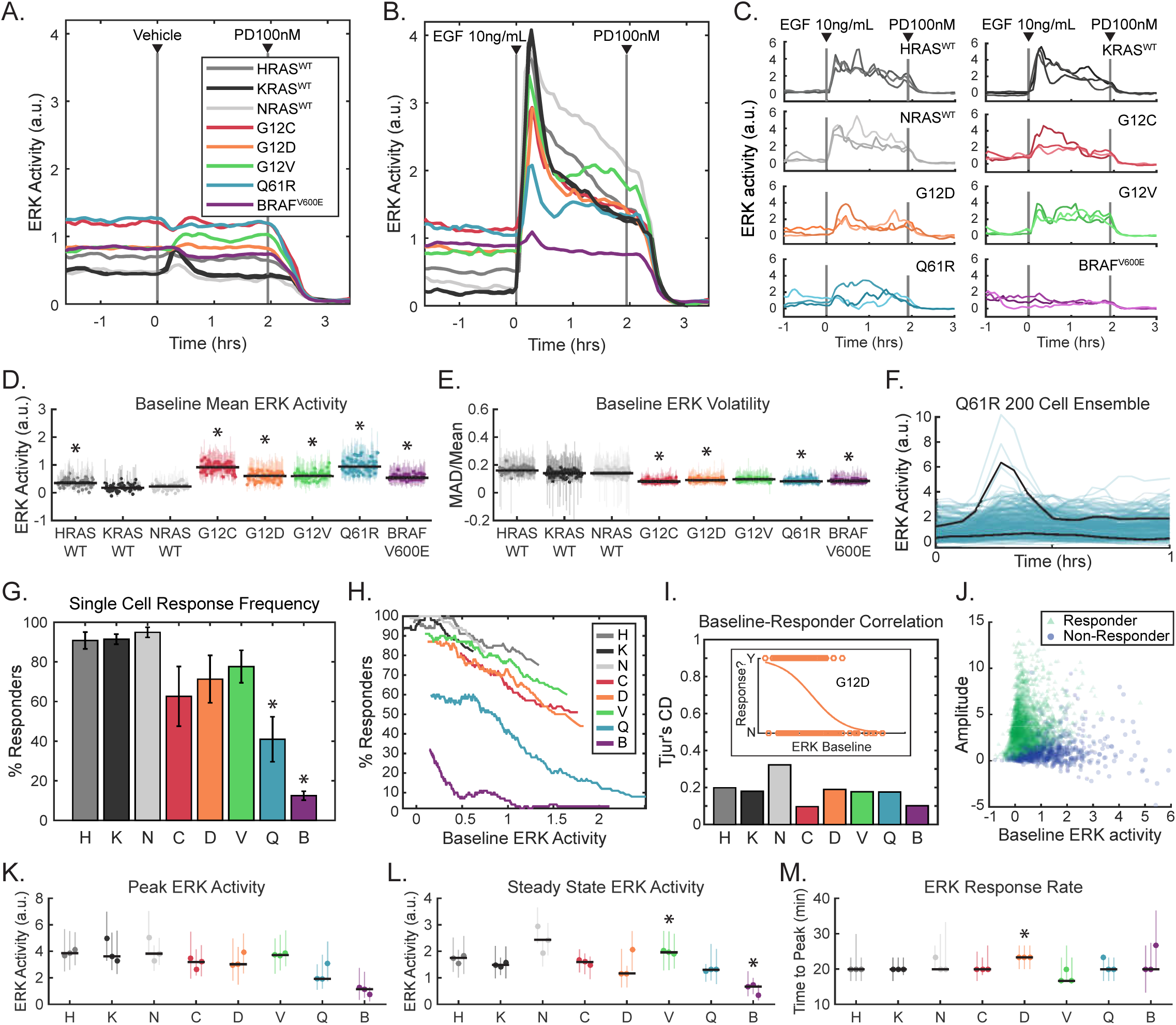
Kinetic analysis of ERK activity for each RAS isoform in response to EGF stimulation. A-E) ERK activity in each of the 8 MEF cell lines, after growth factor withdrawal for 16-24 hours, followed by stimulus consisting of (A) media only, or (B) 10 ng/mL EGF. (A,B) Mean values over 3 replicate cultures. C) Three example single cell traces per cell line from a representative experiment. D) Average baseline (pre-stimulus) ERK activity over 58 replicate cultures per cell line. Each dot represents the mean value across cells in an experiment and lines represent the 25^th^ – 75^th^ percentiles. The black bar denotes the median across all replicates. E) Average volatility (pre-stimulus), reflecting the scale of variation over time, displayed as in D. F-J) Analysis of single-cell response likelihood after EGF stimulus. F) Demonstration of many cells not responding to EGF stimulus in some mutants, in a representative experiment. Black lines highlight one responder and one non-responder cell. G) Likelihood of single cells responding to EGF stimulus, for each cell line, showing mean of 3 replicates with error bars showing one standard deviation. H) Relationship between response likelihood and average baseline ERK activity as a possible correlate, for each cell line. I) Weakness of correlation between baseline ERK activity and response likelihood, measured by Tjur’s coefficient of discrimination (i.e. correlation coefficient for a binary response). Inset shows an example from the KRAS^G12D^ mutant, where dots are scattered per cell by baseline ERK activity (x-axis) and whether that cell responded to EGF (binary y-axis). Orange line indicates the logistic fit. J) Scattered single cell measurements of baseline ERK activity and amplitude of the change after EGF stimulus. Green triangles: cells that responded; blue circles: cells that did not respond. K-M) Analysis of the response to EGF, by filtering to remove cells that do not respond, presented as in D. K) Peak ERK activity reached after EGF stimulus. L) Average ERK activity after 2 hours in the presence of EGF. M) Delay between EGF stimulus and peak ERK activity. Asterisks indicate statistical significance (pFDR < 0.05).

Baseline ERK activity was detectable in all cells but varied in magnitude among the RAS isoforms (Fig. 3D). All KRAS mutant cell lines, as well as HRAS^WT^, exhibited significantly elevated baseline ERK activity compared to KRAS^WT^, with the highest levels observed in KRAS^Q61R^ and KRAS^G12C^. Mutant KRAS and BRAF^V600E^ cells were less volatile over time than KRAS^WT^ cells under baseline conditions (Fig. 3E). Minimal differences in volatility were detected between HRAS^WT^, KRAS^WT^, and NRAS^WT^ cells (Fig. 3E, S2). The higher baseline activity in mutant KRAS isoforms compared to wild type is qualitatively consistent with constitutively higher GTP loading of these GTPase-deficient RAS proteins, which would also reduce variability in RAS activation by obscuring minor spontaneous activation events, such as autocrine signals.

The quantitative parameters of the ERK response to growth factor stimulation, including the amplitude and duration of activity, play an important part in shaping downstream cellular responses (Ebisuya et al., 2005; Nakakuki et al., 2010). We therefore explored the differences between these parameters in mutant and wild type KRAS cells. While the rise and fall of ERK activity occurred with similar kinetics across all KRAS variants, the average peak ERK activity in the KRAS mutant cell lines was unexpectedly equal to or lower than KRAS^WT^ (Fig. 3B). However, differences in the average ERK activity could result from heterogeneity between cells, and upon examination we found that the percentage of cells with a detectable ERK response to EGF was significantly reduced in KRAS mutant lines (Fig. 3F,G). KRAS^Q61R^ cells in particular exhibited drastically reduced response rates. This reduced response could arise from a functionally resistant subpopulation of cells, but could also result from the difficulty of detecting smaller amplitude responses in cells with elevated baseline activity. Therefore, to validate the response measurement, we examined the correlation of response frequency with baseline activity. While the response rate does vary with average baseline activity for most mutants (Fig. 3H), correlation at the single cell level is quite poor for all cell lines (Fig. 3I,J); many high baseline cells clearly respond and many low baseline cells do not. Thus, the population-averaged peak ERK activity is genuinely reduced in KRAS mutant cells by a lower probability of response for each cell.

To remove the bias introduced by non-responding cells and more accurately compare average ERK responses, we filtered the ERK activity dataset to include only cells with a distinguishable response (Fig. 3K-M). In this filtered dataset, the peak ERK responses in KRAS mutant cells were still equivalent to or less than those of the KRAS^WT^ cells (Fig. 3K). Kinetics of the growth factor response also remained similar after correction for non-responding cells (Figure 3L-M). The only distinction observed was that steady state ERK activity 2 hours after stimulation was higher in NRAS^WT^ and KRAS^G12V^ cells, compared with KRAS^WT^, implying a slower attenuation (Fig. 3L). Thus, even accounting for a reduced frequency of response, KRAS mutant cells exhibit peak ERK activity no higher than wild type lines and show no other notable differences. The similarity of peak ERK responses across mutants is unexpected given the range of RAS GTPase activities represented in this panel of cell lines and implies that the upper limit of ERK activity is subject to tight regulation. Altogether, when individual cell variability is accounted for, the only broadly consistent distinction between ERK activity in KRAS^WT^ and mutant isoforms in this system is moderate elevation of unstimulated activity.

### Immediate feedback from ERK is distributed and relatively weak in RAS mutants

The RAS-ERK pathway is subject to multiple feedback effects triggered by ERK activity that could account for the strict moderation of mutant RAS signaling. We therefore assessed the involvement of these feedback loops by comparing activity at several points in the pathway in the absence or presence of the ERK inhibitor SCH772984 (ERKi) (Morris et al., 2013). As an ATP competitive inhibitor with an allosteric mode, ERKi suppresses both the activity of ppERK and the phosphorylation of ERK by MEK (Chaikuad et al., 2014). Consistent with this allosteric inhibition, this treatment had a partial effect on ERK phosphorylation (Fig. 4). To test for ERK-mediated feedback effects, we treated KRAS^WT^, KRAS^G12C^ and KRAS^Q61R^ cells with 100nM ERKi 1 hour prior to EGF stimulation and measured the phosphorylated or active forms of EGFR, RAS, AKT, and MEK by immunoblotting. Indeed, treatment with ERKi resulted in increased MEK dual phosphorylation at Ser217/Ser221 (ppMEK) under both resting and EGF-stimulated conditions in all three cell lines (Fig. 4A,B), confirming the presence of feedback effects on pathway activity upstream of MEK.

**Figure 4.**
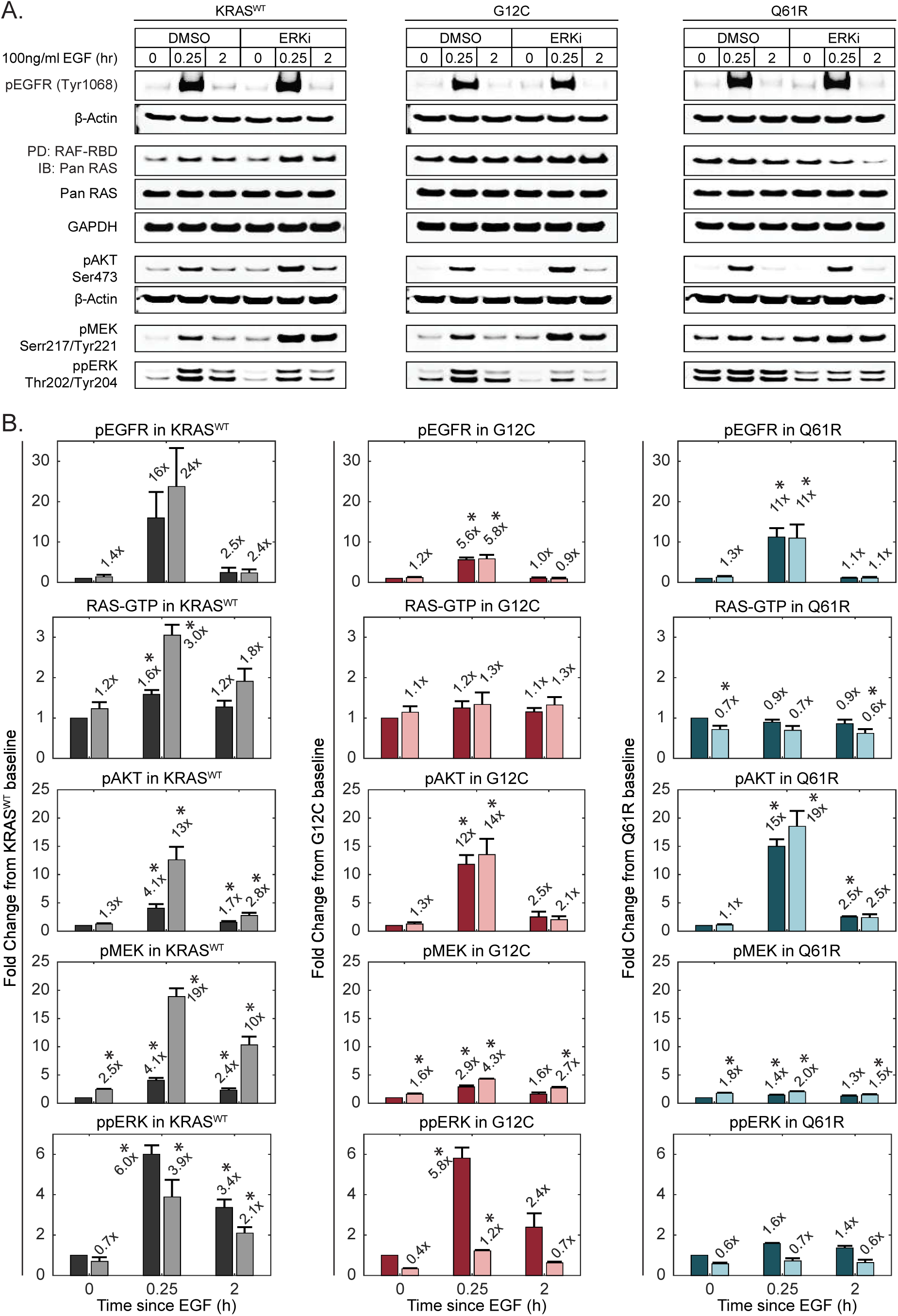
Analysis of ERK-dependent feedback in RAS mutants. A-B) Immunoblot analysis of RAS-ERK pathway activity at multiple levels in the absence or presence of ERK inhibition by 100nM SCH772984 (ERKi). Lysates for the indicated cell lines were collected at baseline, peak (15 min), and steady state (2 hours) time points after treatment with 100 ng/mL EGF. A) Sample blot imagery for each measurement. B) Quantified measurements, shown as fold change relative to the DMSO-treated baseline sample. Values for pEGFR, pAKT, ppMEK, and ppERK were normalized to β-actin; data for RAF-RBD PD/Pan RAS were normalized to total Pan RAS. Bars represent the mean of triplicate measurements and error bars the standard error of the mean. Dark bars: DMSO treated; light bars: ERKi treated. x-axis indicates the duration of EGF treatment. Asterisks indicate statistical significance (p < 0.05).

To explore which steps in the pathway are subject to ERK-mediated feedback, we compared immunoblots of ppMEK, EGFR phosphorylation at Tyr1068 (pEGFR), AKT phosphorylation at Ser473 (pAKT), and RAS activation by pulldown of GTP-bound RAS (RAF-RBD PD; Fig. 4A,B). In unstimulated mutant and wild type KRAS cells, only ppMEK was elevated by ERKi treatment, indicating a significant negative feedback effect due to ERK-mediated inhibitory phosphorylation of RAF (Dougherty et al., 2005). Upon EGF stimulation of KRAS^WT^ cells, we observed the expected increases in pEGFR, pAKT, ppMEK, and RAF-RBD bound RAS. With the exception of pEGFR, all of these species were further increased significantly by ERKi treatment, indicating that under stimulated conditions, ERK-mediated negative feedback also acts at the level of RAS and/or recruitment of GEFs and GAPs, but not receptor activation. These data argue that ERK-mediated negative feedback is distributed throughout the pathway to constrain ERK activation.

However, a different pattern was observed in stimulated KRAS^G12C^ and KRAS^Q61R^ cells. While pEGFR, pAKT, and ppMEK were all significantly increased by EGF stimulation, no significant increase was detected in RAF-RBD pulldown of RAS (Fig. 4A,B). Unlike KRAS^WT^ cells, pEGFR, pAKT, and RAF-RBD-bound RAS were not further increased by ERKi treatment in the KRAS mutant cells. EGF-stimulated ppMEK was significantly increased by ERKi treatment, but with a smaller apparent shift than the corresponding increase in KRAS^WT^ cells. These data suggest weaker ERK-mediated feedback in mutant cells relative to wild type, although they do not rule out the possibility that either the pathway or the assay are saturated under these conditions.

### ERK activity is rescaled bidirectionally, independent of pathway expression levels

The similarity in ERK activity between wild type and mutant KRAS cells contrasts starkly with the conceptual model of RAS mutations hyperactivating the pathway. As this difference is not clearly explained by direct feedback effects from ERK, we employed a mathematical modeling approach to more carefully consider other variables that could account for it (Fig. 5A). In a simple linear view of RAS-to-ERK transduction, the mutant RAS GTPase activities being 50-800 fold lower than wild type would be expected to produce correspondingly large changes in both pre-stimulus and stimulated ERK activity. However, additional variables may modulate the mutant activity. First, variation in expression level of pathway components could compensate for the differences in RAS activity, especially if expression levels become limiting. Second, the affinities of GTP-bound RAS mutants for their effectors are not equivalent to wild type; mutation lowers the affinity for RAF up to 7-fold (Hunter et al., 2015). We therefore designed our modeling approach to predict the expected increase in ERK activity based on known properties of KRAS mutants.

**Figure 5.**
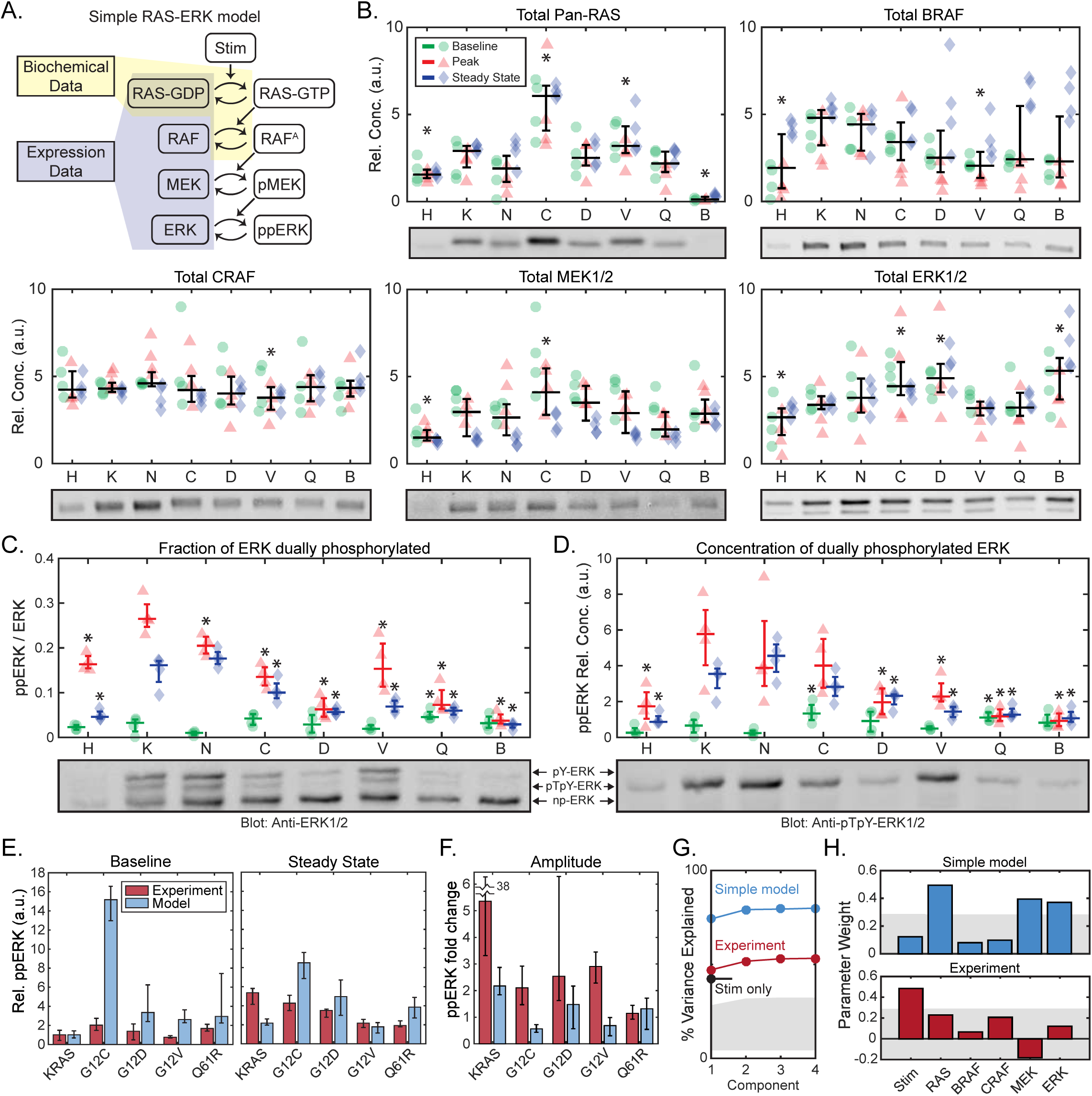
Quantitative analysis of ERK phosphorylation in response to RAS mutation. A) Schematic of a model of the internal factors of the RAS-ERK pathway. Shaded regions indicate portions of the model for which parameter values are available from either published biochemical assays or our immunoblot expression data (B-D). B) Immunoblot measurement of RAS-ERK pathway components in each cell line, at baseline (green circles), peak activity (15 minutes, red triangles) and steady state activity (2 hours, blue diamonds), four replicates each. The median and 25^th^ – 75^th^ percentiles over all conditions are indicated by overlaid whisker plots. Asterisks indicate statistical significance from the KRAS^WT^ cell line. Sample blot imagery is proved underneath each quantification. C-D) Phos-Tag immunoblot measurement of ERK fractional phosphorylation (C) and the relative concentration of dually phosphorylated ERK (D), annotated as in A, but with whisker plots per treatment condition. Sample blot imagery shows anti-ERK1/2 (below C) and anti-ppERK1/2 (below D) for the same blot replicate. E-F) Comparison of the internal RAS-ERK model to experimental data. E) Relative ppERK as predicted by the internal model and measured by immunoblot for the 4 replicates collected, showing the median of the baseline and steady state after EGF treatment (E), and the amplitude of stimulation (F). Error bars show 25^th^-75^th^ percentiles. G-H) Partial least squares regression of both experimental ppERK measurements and predictions via the simple RAS-ERK model. Regression was based on presence/absence of EGF, and expression levels of RAS, BRAF, CRAF, MEK and ERK. G) Percent of variance explained by each PLS model considered, based on how many component terms are allowed. Stim only refers to a PLS model using experimental ppERK data, but only predicting based on the presence/absence of EGF. H) Weights assigned to each parameter in the PLS models. Grey shaded regions indicate the bounds of statistical significance, determined via bootstrapping with scrambled data. Only values that extend beyond the grey regions are statistically significant from zero (p < 0.05).

Our modeling approach requires the relative protein concentration for components of the pathway, which we evaluated by immunoblot at baseline, peak (∼15 minutes), and steady state (∼2 hours) following a 10 ng/mL EGF stimulus (Fig. 5B). In these samples, we also measured the fraction of ERK that is dually-phosphorylated via Phos-Tag immunoblot (Aoki et al., 2013). This measurement of ppERK confirmed the trends observed by FRET measurements: EGF induced a peak in ppERK which then diminished at steady state, and ppERK was elevated in mutant RAS cells only under baseline conditions (Fig. 5C,D). This quantitative dataset rules out two simple explanations for low ERK activity in KRAS mutants. First, compared with KRAS^WT^ cells, total RAS levels were higher in KRAS^G12C^ and KRAS^G12V^ lines, precluding the possibility that the expression level of RAS was compensating for the excess mutant activity (Fig. 5B). Second, levels of BRAF, MEK and ERK, but not CRAF, varied significantly among cell lines, but without a clear pattern or correlation structure. When total expression levels were compared with ERK activity (via EKAR), no significant correlations were found, ruling out the possibility that any individual component acts as a limiting factor. Moreover, dually phosphorylated ERK ranged from only ∼1% to ∼30% of total ERK across all samples (Fig. 5C), confirming that it does not reach saturation.

We parameterized our model using the measured protein levels, along with the biochemical activity of wild type and mutant KRAS proteins previously reported in vitro (Gremer et al., 2011; Hunter et al., 2015; Smith et al., 2013). As the goal of our modeling approach was to identify potential explanations for restrained ERK signaling in KRAS mutant cells, we omitted any additional regulation, such as feedback. Effectively, the model separates operation of the core GTPases and kinases of the cascade, which we term “internal” factors, from “external” factors that that include adapters, scaffolding proteins, and/or phosphatases. By comparing model predictions against experimental data, this approach identifies the differences attributable to external factors. Using a steady state solution of this model, we predicted the baseline and EGF-stimulated steady state levels of ppERK for the isoforms for which biochemical data is available: KRAS^WT^, KRAS^G12C^, KRAS^G12D^, KRAS^G12V^, and KRAS^Q61R^ (Fig. 5E). These simulations confirmed that the experimentally measured ppERK is indeed much lower in KRAS mutants than expected, especially at baseline. Conversely, the amplitude (fold change) in ppERK upon stimulation is greater in the experimental system than in the model, except for KRAS^Q61R^ where differences are indistinguishable (Fig. 5F). This analysis clarifies the role of the external factors in the ERK pathway, revealing that they have a bidirectional effect: they suppress ppERK under both baseline and stimulated conditions, but also amplify the difference between these conditions. The external factors therefore effectively increase the responsiveness of ERK to growth factor stimulation.

To gain further insight into the relative importance of internal and external factors modulating ERK activity, we extended our analysis to test whether ppERK correlates with expression level of pathway components, using partial least square regression (PLSR). We fit both the simulated and measured ppERK against the measured protein expression and the presence of EGF stimulation. PLSR explained 75% of the variance in simulated ppERK, but only 55% of the variance in experimentally measured ppERK (Fig. 5G). In the simulated ppERK data, we found significant correlations with the abundance of RAS, MEK and ERK proteins. In contrast, ppERK in the experimental system was only significantly correlated with the presence of growth factor stimulation (Fig. 5H). Thus, another key function of external factors is to confer robustness to expression level variation in cascade components, extending previous observations that the regulation of ERK phosphorylation is robust to changes in ERK expression level (Fritsche-Guenther et al., 2011). Altogether, our model analysis reveals that external factors increase the dynamic range of ERK response to EGF, and that nearly all of this control lies in mechanisms outside of the linear RAS-to-ERK kinase cascade. This bidirectional effect on ERK activity and the moderate strength of feedback observed in mutant KRAS cells (Fig. 4) imply that ERK activity in the cell is rescaled by mechanisms beyond simple negative feedback (Amit et al., 2007; Dougherty et al., 2005).

### Phosphatases dynamically shape the functional ERK output

Multiple phosphatases are dynamically regulated during growth factor responses (Amit et al., 2007), some directly by ERK (Yoon and Seger, 2006), raising the question of whether such regulation could contribute to the observed rescaling of ERK dynamic range. While phosphatase protein levels can be quantified, phosphatase activity is typically difficult to assay, especially in living cells. However, our experimental system provided a unique opportunity to estimate phosphatase activity acting on ERK substrates by comparing the datasets for ERK activity (measured by FRET, Figs. 2, 3) and the abundance of active ERK molecules (measured by immunoblot, Fig. 4). While these measures are typically considered equivalent under the assumption that phosphatase activity should be stable, our measurements made under identical conditions reveal some differences. For example, we observe significant variation in ppERK across cell lines, while ERK activity is indistinguishable in the same lines, especially at the steady state time point (comparing Fig. 3K,L to Fig. 5D). As noted in our calibration of EKAR, the activity measurement is a ratio of the concentrations of active ERK and any phosphatases that act on ERK substrates. We therefore inferred how this phosphatase activity varied by examining the correspondence between ppERK and activity measurements (Fig. 6A) and estimating the relative phosphatase activity as the ratio of these values (Fig. 6B).

**Figure 6.**
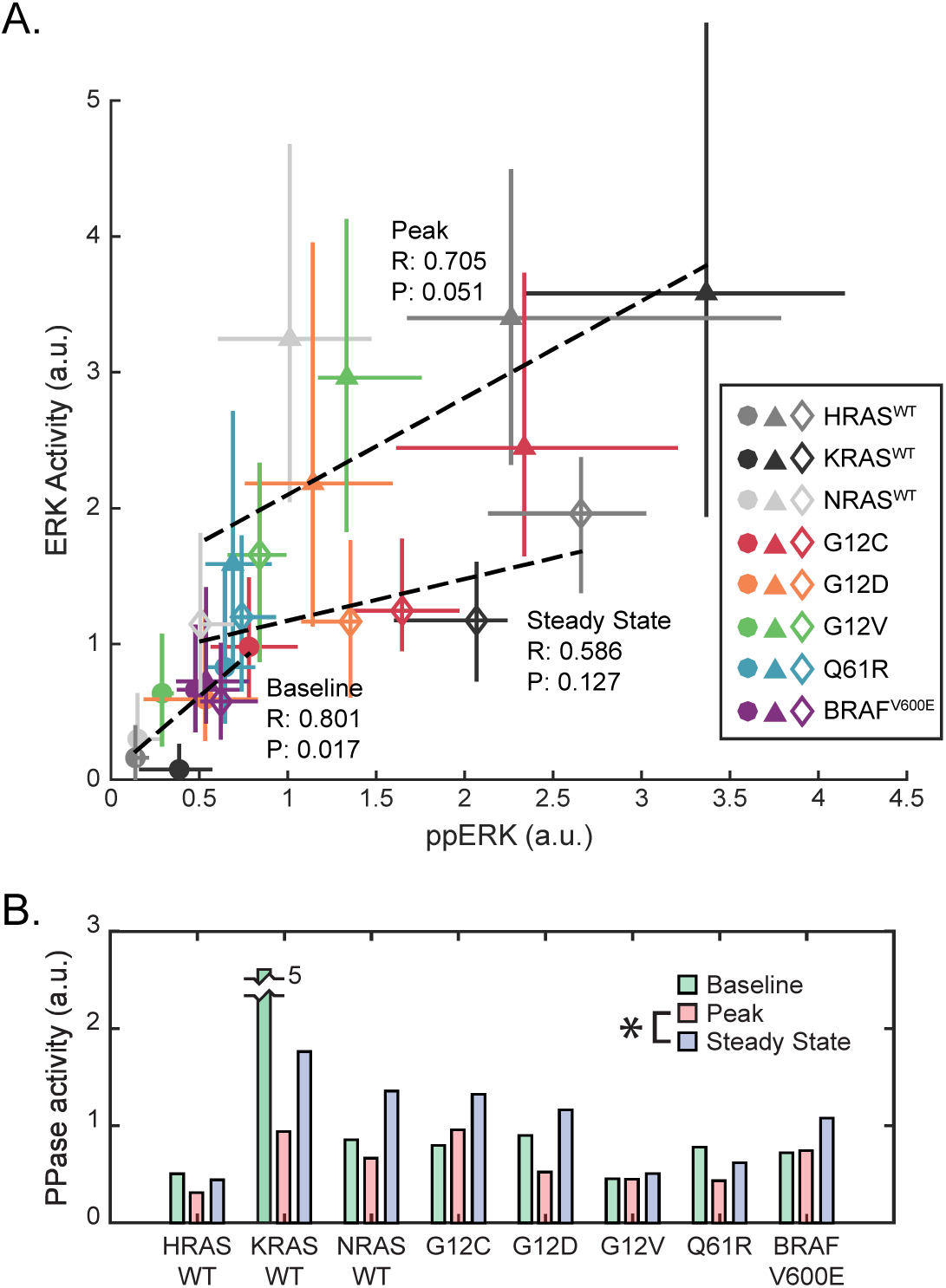
Inference of phosphatase activity on ERK substrates. A) Correlation of ERK activity and ppERK concentration, median of 3 and 4 replicates respectively, per condition. Error bars denote 25^th^-75^th^ percentiles, including single cell distributions for ERK activity. Markers are color-coded by cell line, and marker shape indicates treatment (circle: baseline, triangle: peak, diamond: steady state). Pearson’s correlation coefficients (R) and associated p-values (P) are printed for each treatment. B) Estimate of substrate level phosphatase activity per cell line and treatment, calculated as ppERK/ERK activity. Asterisk indicates significance when comparing all cell lines.

At baseline, phosphatase activity appears nearly uniform across cell lines, except for KRAS^WT^, resulting in a significant linear correlation between ppERK and ERK activity. However, after stimulation, both the slope and correlation are diminished, and variance in the estimated phosphatase activity increases. By the steady state time point, estimated phosphatase activity rises significantly, accounting for 42% - 73% of the drop in ERK activity after 2 hours of stimulation. At this point the different RAS cell lines settle to very similar levels of ERK activity despite varying levels of ppERK. This correlational analysis implies that after stimulation, phosphatase activities and/or levels are regulated in such a manner that they act to normalize the levels of ERK activity, despite residual differences in concentration of ppERK. However, the observation that apparent phosphatase activity is uncorrelated to ppERK at baseline implies dynamic complexity beyond simple regulation by ERK. While this analysis is specific to phosphatases acting at the level of ERK substrates, it is feasible that a similar mechanism is functionally relevant at the level of ERK or MEK, contributing to the bidirectional modulation of ERK activity in cell lines with severe KRAS mutations.

## Discussion

Here, we used a single-cell approach to bring increased temporal resolution and quantitative rigor to the question of how oncogenic RAS mutants alter signaling behavior within the cell. Our analysis provides two major conclusions. First, our systematic dataset reconciles previously conflicting observations of ERK activity driven by RAS mutations. Second, we find that mutant-driven ERK activity is not simply suppressed by negative feedback as previously observed, but instead that multiple mechanisms cooperate to constrain its dynamic range in response to stimuli.

### A unified model of ERK activity as stimulated by growth factors and mutant RAS activity

The canonical view of RAS signaling is that it hyperactivates the ERK/MAPK cascade. Yet, several experimental models of the conversion of a single *Kras* allele from wild type to GTPase-defective mutant have found that this alteration results in no increase, or even a decrease, in activated ERK (Guerra et al., 2003; Huang et al., 2014; Konishi et al., 2007; Tuveson et al., 2004). While similar ERK signaling could result from negative feedback that restrains mutant-driven activity, it is less clear what mechanisms would result in lower ERK activity.

Based on the data presented here, these differences can now be attributed to the temporal and quantitative limitations of the methods previously used to measure ERK activation (primarily uncalibrated ppERK immunoblots). Our dataset recapitulates the reported attributes of mutant KRAS signaling that in isolation appear contradictory: elevated baseline signaling, retained capacity for GF stimulation, and reduced absolute peak upon stimulus. Additionally, we find that RAS-mutant cells have a reduced probability of response that is independent of their current ERK activity, especially in the case of the severe Q61R mutation. This tendency towards unresponsiveness may contribute to the reduced ERK activation observed in RAS mutant cells in the presence of serum or growth factors.

In addition to reconciling previous observations, our analysis also reveals a previously unquantified phenomenon, which is that ERK activity remains unexpectedly responsive to growth factor stimulation in cells carrying mutant KRAS (Fig. 7). This responsiveness arises because, while both baseline and peak GF-stimulated ERK activity are limited in KRAS mutant cells, the relative strength of suppression is greater at baseline than at peak. This observation is consistent with the idea that dynamic range of a signaling pathway is a physiologically important parameter (Janes et al., 2008), and that mechanisms exist to buffer it from deleterious mutations. This effect is only reliably observable through comparisons between isogenic mutant and non-mutant cells using calibrated ERK activation measurements, underscoring the importance of a quantitative, systematic approach to complex signaling networks.

**Figure 7.**
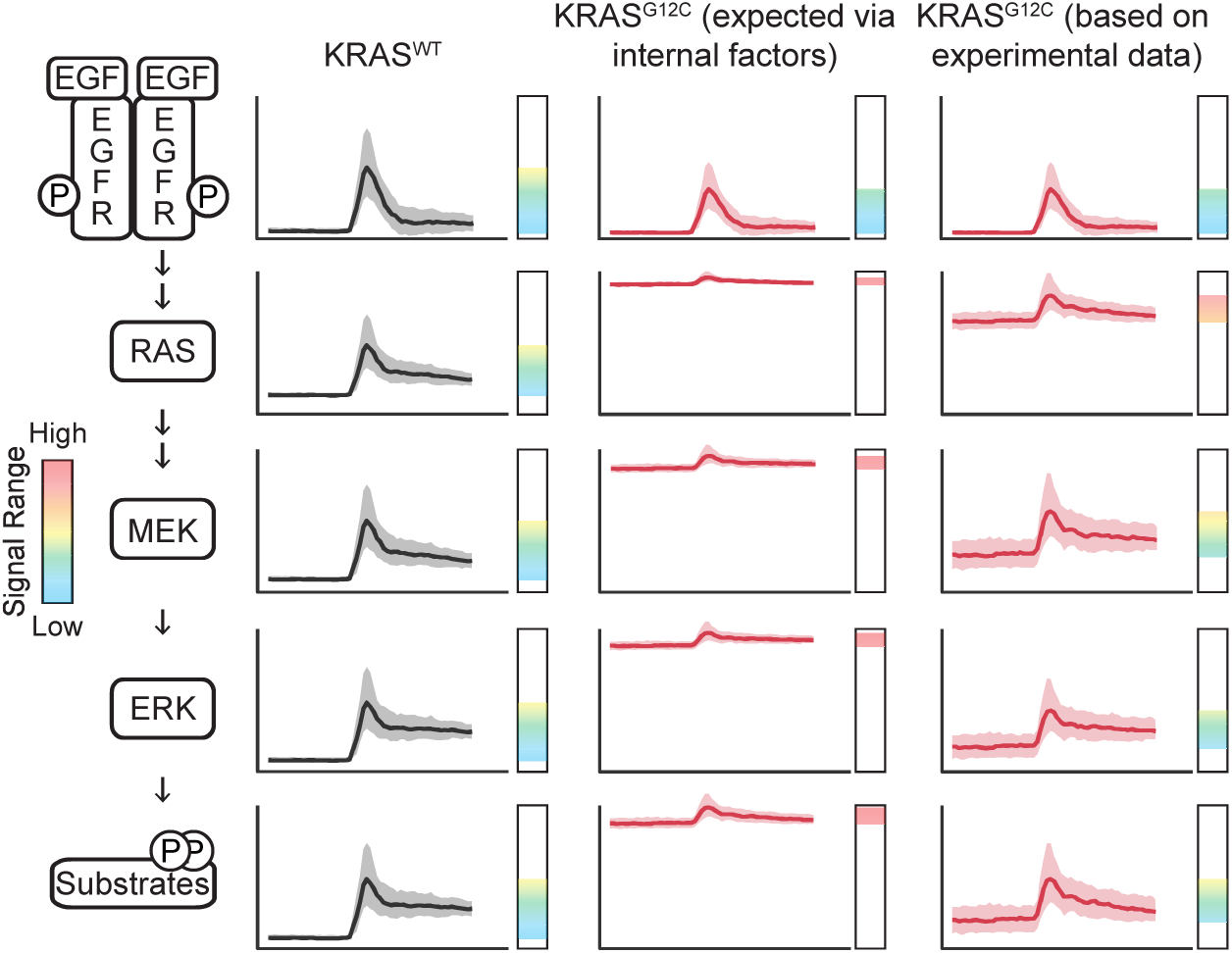
Rescaling and expansion of dynamic range through the RAS/ERK pathway. Diagram depicts the activities we expect at various levels in the RAS/ERK pathway, as time series in our typical experiment, extrapolated from our live-cell data and immunoblot measurements. From left to right, columns depict: the response in KRAS^WT^, the expected response in KRAS^G12C^ cells (as an example KRAS mutant) considering only the internal factors of the RAS-ERK pathway, and observed behavior of KRAS^G12C^ cells. Vertical bars indicate the dynamic range of the activity of that node in the network, with the colored spectrum indicating signal levels from low (blue) to high (yellow - orange) and excessive (red).

### The MAPK pathway as a robust interpreter moderating mutant RAS signaling

The consistency of ERK signaling in the context of RAS mutations or changes in pathway expression has been attributed mainly to negative feedback from ERK (Courtois-Cox et al., 2006; Fritsche-Guenther et al., 2011). However, our systematic analysis reveals a more complex situation with differential suppression distributed across multiple factors. ERK-mediated negative feedback plays a significant role in restraining MEK activation, but that role appears lesser in mutant KRAS cells rather than greater (Fig. 4). At least two models may explain the bidirectional rescaling of ERK signaling that we observe. One possibility is that the RAF/MEK/ERK cascade could act as fold-change detector for variation in EGFR activity (Cohen-Saidon et al., 2009). Fold-change detector models are expected to incorporate motifs such as an incoherent feedforward loop or a non-linear integral feedback loop (Adler et al., 2017). An alternative model is that the pathway is arranged for dose-response alignment (Brent, 2009), which ideally employs push-pull mechanisms or combines negative feedback with a comparator (Andrews et al., 2016). In both models, ERK-mediated negative feedback could play a critical part but would operate in concert with additional regulatory interactions. Such models could explain the complex behavior we observe of restraint of excessive activity from mutant KRAS that also preserves and enhances the ability of the pathway to respond to growth factor stimulus.

Another factor affecting the output of the pathway is the apparent fine-tuning of phosphatase activity acting on ERK substrates. As ERK phosphorylation is often used as a *de facto* measurement of its activity, quantitative effects at the level of substrates have received less attention. Nonetheless, the ability of ERK to maintain phosphorylation of its substrates is inherently limited by the opposing process of dephosphorylation, making this a critical but understudied control point. Our data imply that regulation of this process is significant for an exogenous FRET-based substrate whose sequence is based on the endogenous substrate Cdc25A, warranting further study of this effect on endogenous substrates. This effect could be mediated by direct control of phosphatase activity, or through competition of substrates for the phosphatase (Rowland et al., 2015); future work will be needed to elucidate this mechanism.

Lastly, the potential for each RAS variant line to have been subject to selection during the process of cell line construction and propagation may play a significant role (Li et al., 2018). Cells receiving a RAS insertion that produces sufficiently high levels of expression to drive truly excessive ERK activity could be driven into senescence, and thus prevented from establishing a cell line. Therefore, cells bearing epigenetic modifications or point mutations that moderate the output of ERK could be overrepresented in the surviving population. While our analysis indicates that the expression level of pathway components is insignificant in determining ERK activity in the cell lines assayed, this rules out neither activity-modifying mutations that do not alter expression, nor the existence of an activity threshold above which cells are eliminated by selection. Naturally, the same caveat applies to the vast majority of cell-based experiments on RAS signaling (including transient expression experiments that typically exceed at least one cell cycle). Thus, experimental strategies in which RAS isoforms are abruptly exchanged, and the resulting cellular changes monitored with high temporal resolution, would be informative in understanding the adaptation to a RAS mutation.

### Constraints on RAS-driven signaling in oncogenesis

The ability of the ERK/MAPK pathway to constrain the quantitative effects of mutant KRAS raises important questions for how these mutations function in oncogenesis. In many cancers, RAS mutations are thought to occur very early in oncogenesis, and therefore the homeostatic nature of the pathway likely plays a central role in determining whether a RAS mutant cell progresses toward malignancy (Li et al., 2018). Our data from cells with few other genetic abnormalities can be a considered a model for signaling at this early stage, unique from studies that have investigated mutant RAS in fully developed cancers and focused on treatment of later-stage disease. However, if mutations in RAS lead to only modest changes in ERK signaling, how do they drive progression toward malignancy? One potential model was that excess ERK activity could engage lower-affinity substrates, expanding the effective ERK-driven phosphoproteome to non-traditional targets. However, given the constraints we observe on the magnitude of ERK signaling, it is impractical for these KRAS mutant cells to promote phosphorylation of non-typical ERK targets. Furthermore, KRAS mutant-bearing cells do not show longer duration of peak ERK activity following stimulus than those with KRAS^WT^, so excess activation after growth factor stimulus is also unlikely. Instead, our finding that the over-activating effect of KRAS mutants is limited to chronic baseline elevation implies (1) that chronic moderate signaling is sufficient to drive deleterious phenotypes, and (2) that mutant cells are unlikely to respond to normal low-level signaling.

A strong downstream effect from chronic moderate ERK activity is consistent with current models of some effectors. The ERK target gene Fra-1, a transcription factor whose expression is correlated with cancer invasiveness (Tam et al., 2013), integrates ERK activity over time (Gillies et al 2017). With its slow decay rate (half-life > 5h, (Basbous et al., 2007)), Fra-1 can accumulate to relatively high levels over a long period of moderately elevated ERK activity. Any ERK-induced gene products with similar degradation kinetics will also accumulate over time in cells with baseline ERK elevation. Conversely, gene products subject to rapid degradation kinetics such as c-Fos and Egr-1 would be only weakly elevated in RAS mutants, compared to the large changes in expression driven by sporadic wild type activity. Products under negative regulation, such as those that degrade rapidly even with extended activity (Wilson et al., 2017), may actually be suppressed by the chronic ERK activity. Thus, while enhanced ERK kinase activity as an indicator of early RAS mutant cells is difficult to detect without live-cell measurements, the resulting expression profile - particularly the ratio between long-term and short-term responsive genes - may be more informative.

While the damping of mutant RAS-driven signals at the level of ERK may appear to be a tumor suppressive mechanism, this is not necessarily the case. KRAS mutation frequencies in human cancer and data from mouse models suggest that a limited quantitative range of RAS signal (a “sweet spot”) is critical for the development of tumors (Li et al., 2018; Sarkisian et al., 2007). Pathway constraints could help RAS mutant cells to stay within this range and evade senescence or cell death due to excessive ERK activation. This paradox raises the question of whether RAS mutations are more common than downstream mutations (such as MEK or ERK) in cancer and related syndromes such as RASopathies because they are strong enough to induce increased ERK activity, or rather because they are more constrained and able to escape selection by senescence.

## Acknowledgements

Funding for this work was provided by the National Institute of General Medical Sciences (1R01GM115650 to JGA), the Department of Defense Neurofibromatosis Research Program (W81XWH-16-1-0085 to JGA), and the National Cancer Institute (K01CA197138 to JMS and 1R35CA197709 to FM). Flow-cytometry services were supported by the UC Davis Comprehensive Cancer Center Support Grant (CCSG) awarded by the National Cancer Institute (NCI P30CA093373), and we acknowledge the expert cell sorting assistance of Dr. Bridget McLaughlin and Jonathan Van Dyke. All cell lines were kindly provided by Dom Esposito at the National Cancer Institute Ras Initiative, Frederick, MD.

## Author Contributions

Conceptualized the study: MP, JGA, FM

Designed the experiments: TEG, MP, JS, FM, JGA

Prepared cell lines: TEG, CT, JMS, FM

Performed live-cell experiments: TEG, CT

Performed western blot experiments: TEG, MP, JMS

Prepared mathematical models: MP

Analyzed the data: TEG, MP, JS, JGA

Wrote the manuscript: TEG, MP, JGA

Reviewed/edited the manuscript: TEG, MP, JMS, CT, FM, JGA

## Materials and Methods

### Cell Culture

Mouse embryonic fibroblasts expressing a single RAS isoform were obtained from the National Cancer Institute, Frederick, MD. Cells were cultured in DMEM supplemented with 0.2% bovine serum albumin (BSA) and 2.5 µg/mL puromycin or 4 µg/mL blasticidin. For imaging experiments, cells were cultured in a custom imaging media composed of DMEM lacking phenol red, folate and riboflavin, glucose, glutamine, and pyruvate, supplemented with 0.1% BSA, 4mM L-glutamine, and 25mM glucose.

### Reporter Cell Line Construction

Cells were electroporated using a Lonza Nucleofector electroporator. EKAR3 was stably integrated into cells using the piggyBAC transposase system (Pargett et al., 2017). Positive integrants were selected by fluorescence-based cell sorting.

### Live Cell Microscopy

Multi-well plates with #1.5 glass bottoms were coated with collagen and seeded with reporter cell lines one day prior to imaging. Prepared culture plates were imaged on a Nikon Ti-E inverted microscope with a stage-top incubator to maintain the culture at 37°C and 5% CO2 throughout the experiment. Microscopy and image processing performed as described in (Pargett et al, 2017). Imaging sites within each well were selected and imaged sequentially at each acquisition time, automated via the NIS-Elements AR software. Images were captured using a 20x/0.75 NA objective and an Andor Zyla 5.5 scMOS camera.

### Immunofluorescence Microscopy

After growth and treatment as indicated on glass-bottom 96-well plates, cells were fixed for 30 min at room temperature with a freshly prepared solution of 12% paraformaldehyde in PBS and permeabilized with 1% Triton X-100. Samples were then stained with primary and secondary antibodies in PBS+0.1% Triton X-100+2% bovine serum albumin, and images were captured on a Nikon Ti-E inverted microscope with a 20x/0.75 NA objective with an Andor Zyla 5.5 scMOS camera.

### Image Processing

Imaging data were processed to segment and average pixels within each identified cell’s nucleus and cytoplasm, using a custom procedure written for MATLAB (Pargett et al., 2017). The procedure accessed image data from ND2 files generated by NIS Elements, using the Bio-Formats MATLAB toolbox, and tracked single cell positions over time using uTrack 2.0 (Jaqaman et al., 2008). The resulting single cell time series traces were filtered for quality (minimum length of trace, maximum number of contiguous missing or corrupt data points), and ratiometric reporter levels calculated. EKAR3 level was calculated as 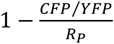, where *CFP* and *YFP* are the pixel intensities of the cyan and yellow channels, respectively, and *R_P_* is the ratio of total power collected in cyan over that of yellow (each computed as the spectral products of relative excitation intensity, exposure time, molar extinction coefficient, quantum yield, light source spectrum, filter transmissivities, and fluorophore absorption and emission spectra). See Supplemental Methods for detailed interpretation of the EKAR3 signal.

### Immunoblotting

For immunoblot experiments assaying pathway activity and feedback sensitivity (Figure 4), cells were seeded at a density of 2.5×10^6^ cells per 10 cm plate and starved of growth factor for 6 hours in imaging media. Cells were pre-treated with DMSO or 100nM SCH772983 (Selleckchem) (Morris et al., 2013) for the last hour of starvation. Cells were then stimulated with vehicle or 100ng/ml EGF for 15 minutes or 2 hours and lysed with Cell Lysis Buffer (50mM Tris pH 7.5, 10mM MgCl2, 0.5M NaCl, 2% Igepal) (Cytoskeleton Inc.) containing protease and phoshatase inhibitors (Pierce/Thermo Fisher Scientific). Lysates were clarified by centrifugation at 10,000 rpm for 2 minutes at 4°C and snap frozen in liquid nitrogen with protein concentrations measured using the BCA protein assay (Pierce/Thermo Fisher Scientific). For RAS activation assays, 300µg of total cell protein was used to pull-down GTP-bound RAS/RAF-RBD complexes according to the manufacturer’s instructions (Cytoskeleton). Activated RAS or 20µg of total cell protein were separated using NuPAGE Novex Bis-Tris gels (Invitrogen/Thermo Fisher Scientific) and transferred to PVDF membrane using an iBlot™ 2 dry blotting system (Invitrogen/Thermo Fisher Scientific). Immunoblot data for this assay were analyzed using the Odyssey® application software v3.030 as described previously (Silva and McMahon, 2014). Statistical significance was determined by t-test analyses of three independent experiments.

For immunoblot analysis of pathway expression levels (Figure 5), we prepared 4 replicate samples of each treatment (baseline, stimulation, or steady state) for each of the 8 cell lines. Cells were plated in 10cm plates at a density equivalent to that in our imaging experiments (1.7 x 10^6^ cells, ∼30,000 cells/cm2). Cultures to be stimulated were treated with EGF at 10ng/mL EGF for 15 minutes or 2 hours, matching the timing of peak activity in live cell data. Cultures were lysed in 500 uL RIPA buffer with protease and phosphatase inhibitors and clarified as above. These lysates were assayed for protein content by DC protein assay (Bio-Rad) and frozen. Immunoblotting was performed using SDS-PAGE with 12.5% acrylamide or 4-15% gradient gels (Bio-Rad, Cat #4561046, Cat #4561086).

To include the many samples in this dataset on the same scale via immunoblot, we employed lane-to-lane normalization by total protein load and blot-to-blot normalization by including one sample, the “control”, from each blot together on a reference blot. In this way, variations in staining efficiency among membranes are accounted for by scaling all lanes (per target protein) such that the control sample matches its intensity on the reference blot. Normalization to protein load is performed by Ponceau S stain, and is applied for all samples prior to normalization across blots, such that variation in the Ponceau stain is also included in the reference blot. Intensity measurements were performed using ImageJ, with background samples collected adjacent to each band/region of interest.

### Phos-Tag immunoblotting

We employed the Phos-Tag method using precast gels (SuperSep Phos-Tag 12.5% Cat #195-17991, and 7.5% Cat #192-18001, FUJIFILM Wako Chemicals). However, in accordance with previous observations with these gels (Kinoshita-Kikuta et al., 2012) and the observation that they likely have excess Phos-Tag reagent, we use samples collected in EDTA-containing RIPA buffer and we perform electrophoresis with Tris-Glycine running buffer (as with above). Phos-Tag fractional phosphorylation measurements are internally controlled and required neither cross-load or cross-blot normalization. See Appendix for detailed supplemental methods.

### Statistical Analysis

For all imaging experiments shown, a minimum of 100 cells were imaged and tracked for each condition. Single-cell data points were excluded as outliers if greater than six standard deviations from the dataset mean. For all analyses, at least three independent experimental replicates were performed. Where indicated, single cell data were normalized to the median value of the PD0325901-treated period. All statistical and computational tasks were performed using MATLAB. Each single cell trace was normalized to the minimum value in a 1 hour window following treatment with 100nM PD0325901. Baseline values were calculated by taking the mean of the 2 hour window prior to stimulation for each cell. The mean was calculated from a 2 hour window following treatment with the specified growth factor or vehicle control. Volatility is calculated as the scaled mean absolute derivative, i.e. the sum of the absolute value of the derivative over a 2 hour window following stimulation, divided by the mean of the same window. Responders were defined as cells with: (1) post-stimulation ERK activity that significantly increased at least 5% compared to the average baseline value, and (2) a higher maximum derivative at the time of stimulation compared to the baseline region. Response rate was defined as the time from stimulation to the peak ERK activity in each responding cell.

Unless otherwise indicated, each statistical comparison was made by t-test with unequal variances, and false discovery rate was controlled within each dataset via the Benjamini and Hochberg Step-Up procedure (α = 0.05). Where replicates were available at the single cell level as well as across experiments, the variance of the mean for each experiment was determined from single-cell samples and added to variance across experiments. This corresponds to a linear error model: *ε_i_* = *ε_cell_* + *ε_exp_*, where there error (from the mean) of an individual cell *ε_i_* equals the sum of the errors arising from cell-to-cell variation *ε_cell_* and from experiment variation *ε_exp_*.

